# Exocyst Inactivation in Urothelial Cells Disrupts Autophagy and Activates non-canonical NF-κB

**DOI:** 10.1101/2021.07.05.451173

**Authors:** Michael A. Ortega, Ross K. Villiger, Malia Harrison-Chau, Suzanna Lieu, Kadee-Kalia Tamashiro, Amanda J. Lee, Brent A. Fujimoto, Geetika Y. Patwardhan, Joshua Kepler, Ben Fogelgren

**Author notes:** Corresponding Author: Ben Fogelgren.

## Abstract

Ureter obstruction is a highly prevalent event during embryonic development and is a major cause of pediatric kidney disease. We have reported that ureteric bud-specific ablation of the exocyst Exoc5 subunit in late-murine gestation results in failure of urothelial stratification, cell death, and complete ureter obstruction. However, the mechanistic connection between disrupted exocyst activity, urothelial cell death, and subsequent ureter obstruction was unclear. Here, we report that inhibited urothelial stratification does not drive cell death during ureter development. Instead, we demonstrate that the exocyst plays a critical role in autophagy in urothelial cells, and that disruption of autophagy activates a urothelial NF-κB stress response. Impaired autophagy first provokes canonical NF-κB activity which is progressively followed by increasing non-canonical NF-κB activity and cell death if the stress remains unresolved. Furthermore, we demonstrate that ureter obstructions can be completely rescued in Exoc5 conditional knockout mice by administering a single dose of pan-caspase inhibitor z-VAD-FMK at E16.5 prior to urothelial cell death. Taken together, ablation of Exoc5 disrupts autophagic stress response and activates progressive NF-κB signaling which promotes obstructive uropathy.

## Introduction

Ureter obstruction during fetal development is a common cause of children being born with congenital anomalies of the kidney and urinary tract (CAKUT) (Chua *et al*, 2019; Johansen *et al*, 2021; Verbitsky *et al*, 2019). The most prevalent site of obstruction occurs at the ureteropelvic junction (UPJ), where the renal pelvis transitions into the upper ureter, resulting in restricted urine flow that can cause lasting kidney damage (Roth *et al*, 2002). Whether the obstruction resolves naturally or is surgically corrected, as many as 70% of patients with congenital obstructive uropathy (COU) will develop a gradual loss of kidney function and progress to end-stage renal disease by the age of 20 (Chevalier *et al*, 2010; Craven *et al*, 2007; Mesrobian & Mirza, 2012; Miklovicova *et al*, 2008). However, there remains a limited understanding of the mechanisms that govern the embryonic onset of COU.

Several mouse models have implicated urothelial abnormalities as a major driver behind ureter obstructions (Jackson *et al*, 2020). The urothelium is a specialized stratified epithelium that functions as a urine permeability barrier along the upper urinary tract and bladder. We have previously reported that conditional knock out of the exocyst *Exoc5* gene in ureteric bud cells disrupts the urothelial stratification process in embryonic ureters, which subsequently triggers cell death between E16.5-E17.5 (Fogelgren *et al*, 2015). The *Exoc5* conditional knockout was achieved using the Ksp-cadherin Cre mouse strain (Cre^Ksp^), which we demonstrated to be active <E12.5, well before the urothelial cell death began, indicating the disruption of stratification may be key to triggering cell death. The wave of cell death in *Exoc5*^*FL/FL*^*;Cre*^*Ksp*^ ureter is followed by a wound healing response that causes UPJ lumen obliteration through the activation and expanse of myofibroblasts (Fogelgren *et al*., 2015; Lee *et al*, 2016). The resulting phenotype of the *Exoc5*^*FL/FL*^*;Cre*^*Ksp*^ mouse (Exoc5-CKO) is similar to intrinsic human congenital UPJ obstructions and is a valuable model for investigating the underlying mechanisms of ureter obstruction.

EXOC5 is a core component of the highly-conserved octameric exocyst protein complex (consisting of EXOC1-8) that mediates the targeting and docking of intracellular vesicles (Ahmed *et al*, 2018; Heider *et al*, 2016; Mei *et al*, 2018; Morin *et al*, 2010; TerBush *et al*, 1996). Since the exocyst associates with specific vesicles through members of the Rab GTPases family, the exocyst is often classified as a Rab effector complex. The localization and assembly of the exocyst holocomplex is guided in part by members of the Ras superfamily of small GTPases such as RALA and RALB, which regulate distinct biological processes by interacting with different exocyst subunits. As an example, the RALA-EXOC2 complex mediates polarity by trafficking proteins to the basolateral membrane in epithelial cells (Moskalenko *et al*, 2002), while the RALB-EXOC2 complex can directly activate innate immunity response and restrict apoptosis by engaging the IkB kinase family member TBK1 (Chien *et al*, 2006; Shipitsin & Feig, 2004). Alternatively, RALB-EXOC8 can promote autophagy by acting as an assembly scaffold for ULK1 and Beclin1-VPS34 during nutrient deprivation (Bodemann *et al*, 2011; Martin *et al*, 2014; Singh *et al*, 2019). Since these different interacting combinations can activate distinct responses, a clearer understanding of the role the exocyst complex performs in cellular stress response is necessary (Simicek *et al*, 2013).

For example, studies in *Drosophila* have revealed that exocyst-mediated autophagy is tissue-specific and context dependent (Mohseni *et al*, 2009; Tracy *et al*, 2016). Autophagy is an evolutionarily conserved lysosomal degradation process known to be highly active during differentiation and development (Mizushima & Levine, 2010), and is critical for many physiological events, such as responding to cellular stress by maintaining homeostasis through the clearance of damaged organelles and proteins (Kroemer *et al*, 2010; Tang *et al*, 2020). Interestingly, while autophagy appears to be dispensable for mammalian ureter and kidney development (Goodall *et al*, 2016; Gump *et al*, 2014; Komatsu *et al*, 2005; Kuma *et al*, 2004; Nezis *et al*, 2010; Thorburn *et al*, 2014), several mouse models utilizing tissue-specific autophagy related gene (ATG) knockout demonstrate that deficiencies in autophagy promote progressive pathology by limiting the organism’s ability to respond to stress (Bechtel *et al*, 2013; Hartleben *et al*, 2010; Kim *et al*, 2012; Takahashi *et al*, 2012).

Here, we test if ureter obstruction in *Exoc5* CKO mice is the result of failed urothelial stratification or because of the unresolved urothelial cell death, and whether prevention of urothelial differentiation during ureter development directly causes to cell death. Furthermore, we used a ureter explant model to determine if the fibroproliferative wound healing reaction that obliterates the ureter lumen is activated in the absence of urine. Our data show that autophagy is impaired after urothelial *Exoc5* ablation, which triggers an increasing prevalence of the TNF superfamily receptor Fn14, a potent activator of the noncanonical NF-κB pathway. Our data demonstrate that inhibiting autophagy in urothelial cells provokes an initial p65 canonical NF-κB response that is progressively followed by a secondary p52 non-canonical NF-κB response and then cell death. Lastly, we investigate if the ureter urothelial cell death is critical to COU in our *Exoc5* CKO mice through rescue experiments with pan-caspase z-VAD-fmk inhibitor at E16.5. Taken together, our data demonstrate impaired exocyst-mediated autophagy in urothelial cells activates progressive NF-κB signaling that enhanced cell death and promotes the onset of COU.

## Results

### Differentiating urothelial cells initiate cell death independent of failed stratification

Abnormal urothelial differentiation has been implicated in mouse models and patients with CAKUT. To investigate the underlying cause of the obstruction in *Exoc5*^*FL/FL*^*;Ksp-Cre* (CKO) embryonic ureters, we developed an *ex vivo* ureter explant culture system. Wild type C57BL/6J mouse embryonic ureters were collected at E15.5 and cultured for 72 hours on semi-permeable supports at the air-liquid interface (Figure 1A). In this system, the explant urothelium starts as a single epithelial monolayer, as shown by immunostaining for E-cadherin (green) and smooth muscle actin (SMA, red) (E15.5, t=0), and successfully differentiates into a stratified epithelium with uroplakin staining on the luminal surface (E18.5, t=72hrs). Uroplakins are produced by superficial cells, thus indicating successful urothelial differentiation in an *ex vivo* setting over a time course similar to *in vivo* ureter development. After 72 hours, the explant ureters showed increased overall growth and continued to have regular peristalsis, indicating the successful maturation of the smooth muscle layer that surrounds the ureters (Figure 1B).

**Figure 1.**
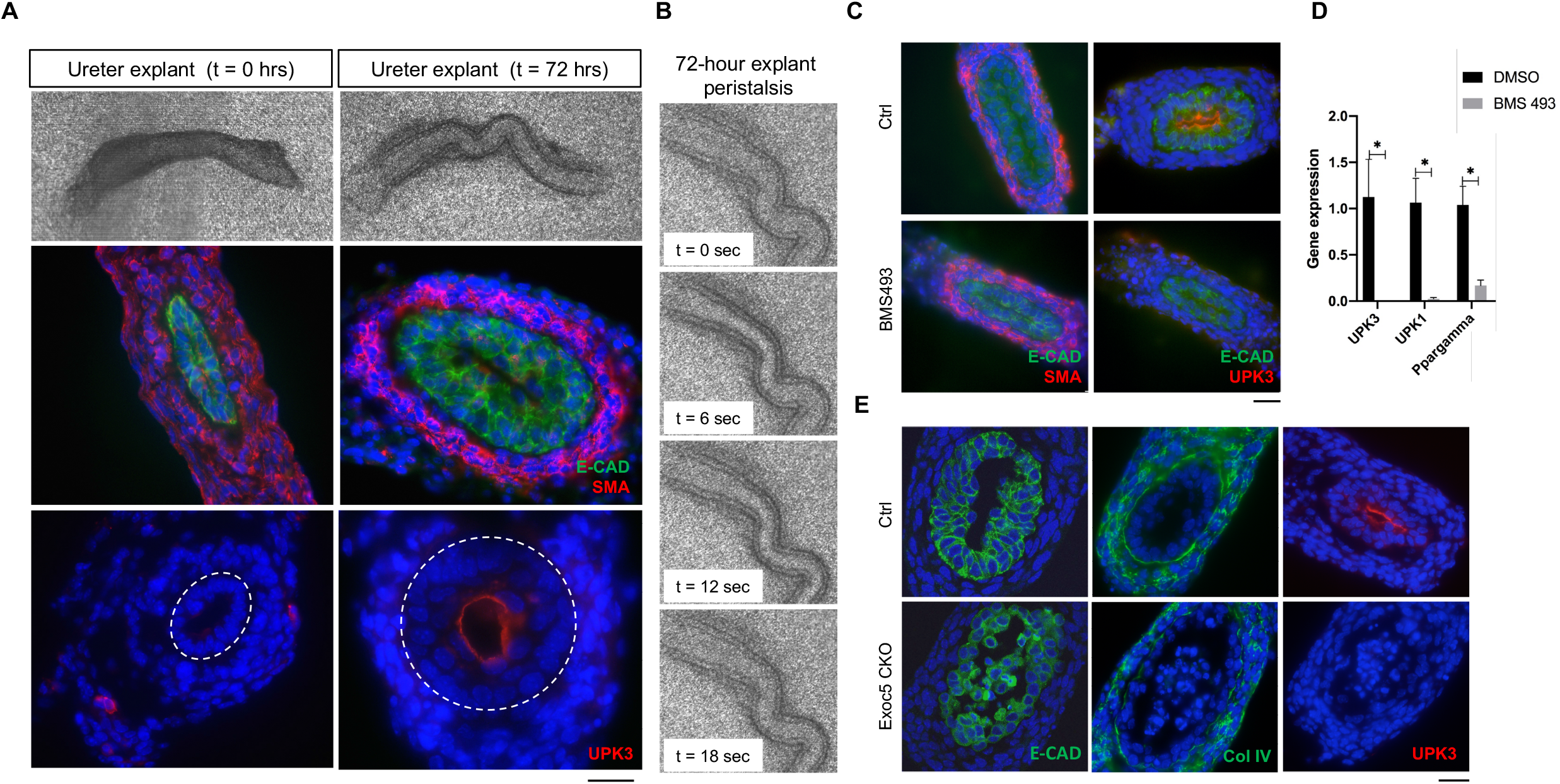
Urothelial cell death occurs independent of failed stratification. (A) *Ex vivo* ureter explant culture model results in viable *Exoc5*^*FL/FL*^ (control) ureter explants after 72 hours. Growth of E15.5 (t=0 hours) control ureter explants shown after cultured for 72 hours. Explants were fixed, cryosectioned, and immunostained for E-cadherin (green) and SMA (red), which revealed urothelial stratification and strengthening of the smooth muscle layer. Staining for uroplakin-3 (UPK3) (red) demonstrated successful differentiation of superficial cells *ex vivo*. (B) Time lapse images of ureter explant peristalsis confirmed functional viability of ureters after 72 hours. (C) Wildtype ureter *ex vivo* treated with retinoic acid inhibitor BMS493 at 100 μM for 72 hours developed a monolayered epithelium and lacked uroplakin expression with no apparent cell death, while vehicle control treated ureter explants developed a multilayered urothelium as indicated by E-cadherin (green) and uroplakin-3 (red). (D) Real-time qPCR confirmed uroplakin-3 (UPK3), uroplakin-1 (UPK1), and PPARγ (PPARgamma) downregulation in BMS493-treated ureter explants. Statistical significance denoted by p* ≤ 0.05. (E) *Exoc5* CKO *ex vivo* ureter explants cultured for 72h hours displayed a disrupted urothelial layer with epithelial cells detaching from the smooth muscle layer and entering the lumen. *Exoc5*^*FL/FL*^ control *ex vivo* ureter cultures showed differentiation of a multilayered urothelium indicated by E-cadherin (green) and presence of uroplakin-3 (red). Scale bars = 20 µM.

Upon establishing this *ex vivo* ureter model, we wanted to determine if failed differentiation was the underlying cause of urothelial cell death observed in *Exoc5* CKO mice. To test this, we treated E15.5 explant ureters with either vehicle control or retinoic acid receptor inverse agonist BMS 493 for 72 hours, since retinoic acid signaling is known to be necessary for urothelial differentiation (Gandhi *et al*, 2013). Explant ureters treated with BMS 493 were unable to differentiate a multilayered epithelium, as shown with E-cadherin/SMA immunohistochemistry (Figure 1C). The BMS 493-treated ureters also completely lacked uroplakin expression (Figure 1D), however, no signs of TUNEL positive urothelial cell death (data not shown) or lumen obstruction were detected. Real-time qPCR analysis of BMS 493-treated samples showed Pparγ, a downstream target of RA, was also strongly down regulated, confirming that BMS 493 successfully disrupted the RA signaling pathway and urothelial differentiation (Figure 1D). Furthermore, EXOC5 concentration at the apical luminal membrane of urothelial cells mirrored previous observations with in vivo Exoc5 CKO ureters which remain unchanged with BMS 493 treatment (Figure EV1) (Lee et al., 2016).

Next, we wanted to use this ureter explant model to determine if the *Exoc5* CKO urothelial cells still underwent cell death, and if the fibroproliferative obstruction would be induced, in the absence of urine flow. For this, we cultured *Exoc5* CKO and littermate control E15.5 explants for 72 hours, followed by immunohistochemistry. As with the wild type ureter explants, control ureter explants showed the presence of a normal multilayered urothelium after 72 hours, while CKO ureters displayed a disrupted urothelium with urothelial cells sloughing off and entering the luminal space as seen with E-cadherin (green) (Figure 1E). As described *in vivo* (Fogelgren *et al*., 2015; Lee *et al*., 2016), *Exoc5* CKO ureter explants also showed no uroplakin expression, indicating the urothelial progenitors failed to differentiate into superficial cells. However, unlike the pathology of *Exoc5* CKO ureters *in vivo*, the distribution of collagen IV indicated an intact basement membrane, with no expansion of mesenchymal cells, suggesting a fibroproliferative response was not activated as a result of the urothelial cell death. Furthermore, we did not find a difference in proliferation as measured by Ki67 (Figure EV2) and only cells that had sloughed off from the basement membrane were found to be cleaved caspase 3 positive (Figure EV3). Taken together, these results suggest that the epithelial cells comprising the urothelial progenitor monolayer initiate cell death independently of failed differentiation or the presence of urine.

### NF-κB activator Fn14 is highly upregulated in *Exoc5* CKO ureters

We found that only a small number of cells that have sloughed off from the basement membrane of Exoc5 CKO ureters are TUNEL positive (Figure EV4). To identify molecules potentially involved in the urothelial cell death event in *Exoc5* CKO ureters, we performed gene profiling on microdissected E16.5 ureter RNA samples using Affymetrix Clariom D GeneChip microarrays. As expected based on our previously reported data (Fogelgren *et al*., 2015), we measured strongly decreased expression of uroplakin genes in *Exoc5* CKO ureters (Figure 2A). One of the most upregulated individual genes was *Fn14* (*Tnfrsf12a*) which is a TNF superfamily receptor known to play a role in both canonical and non-canonical NF-κB signaling (Figure 2A). Fn14 has only one known ligand, a cytokine named TWEAK (tumor necrosis factor-like weak inducer of apoptosis), and has been shown to be a stress-response gene in many tissues (Muñoz-García *et al*, 2006; Nagy *et al*, 2021; Peng *et al*, 2018; Unudurthi *et al*, 2020; Zhao *et al*, 2007). *Fn14* is often strongly upregulated after cell damage or oncogenic transformation (Johnston *et al*, 2015), however there are no published reports of its activity in urothelial cells. We then performed KEGG pathway analysis which implicated NF-κB signaling among the most upregulated pathways in response to *Exoc5* ablation (Figure 2B). qPCR validation confirmed the loss of uroplakin expression in these *Exoc5* CKO samples and that expression of *Fn14* and its ligand *TWEAK* were both increased by more than 30 fold at E16.5 (Figure 2C). Since Fn14 has a relatively short half-life of ∼74min (Gurunathan *et al*, 2014), we performed western blotting on isolated E16.5 and at E17.5 ureters to measure protein levels. Fn14 maintained relatively low protein levels at E16.5, however, there was a strong buildup of Fn14 in E17.5 *Exoc5* CKO ureters, indicating that Fn14 was elevated during this period of stress (Figure 2D). The degree of Fn14 increase in E17.5 *Exoc5* CKO ureters varied but was consistently and significantly higher compared to other genotypes (Figure 2E). In order to determine where Fn14 was being expressed, we performed immunohistochemistry and identified Fn14 was strongly upregulated in the urothelium, but also in the underlying mesenchymal cells (Figure 2F). These data demonstrate that the NF-κB signaling activator Fn14 robustly responds to stress induced by *Exoc5* CKO during ureter development.

**Figure 2.**
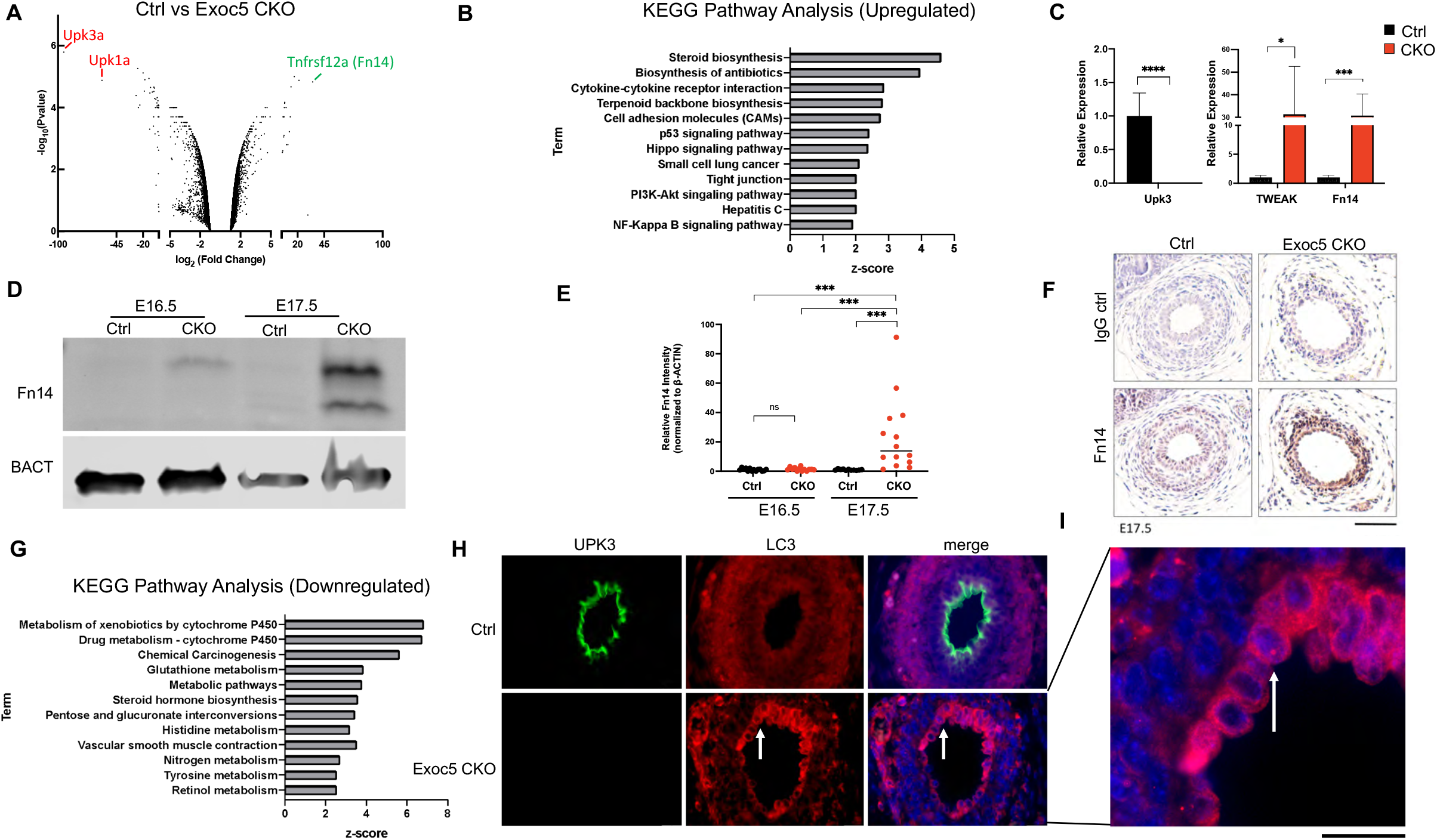
NF-kB signaling and autophagy are significantly impacted by *Exoc5* ablation in developing mouse ureter. (A) Gene expression profiling of total ureter RNA from control and *Exoc5* CKO mice at E16.5 (n=3 each group). Volcano plot displaying expression analysis shows uroplakin genes as significantly downregulated (red) and NF-kB signaling mediator, TNFRSF12A (Fn14), as significantly upregulated (green). (B) NF-kB signaling was among the most upregulated pathways identified by KEGG Pathway Analysis. (C) Real-time qPCR confirmed uroplakin-3 downregulation and Fn14 upregulation in *Exoc5* CKO E16.5 ureters. In addition, the cytokine TWEAK was significantly upregulated along with its receptor Fn14. (D) Western blot analysis of *Exoc5* CKO ureters showed robust expression of Fn14 at E17.5 when compared to control ureters or even *Exoc5* at E16.5. (E) Quantification of Fn14 detected by western blots in E16.5 and E17.5 control and *Exoc5* CKO ureters (n=14,14,11,14, respectively) shown with each dot representing a mouse. (F) Immunohistochemistry of E17.5 control and E*xoc5* CKO ureters demonstrated Fn14 localization to the urothelial cells surrounding the ureter lumen. (G) KEGG Pathway Analysis of the most downregulated pathways indicated broad metabolic disruption. (H) Immunohistochemistry of E17.5 control and E*xoc5* CKO ureters displayed differences in LC3 distribution and onset of LC3 puncta formation (red, arrow). As expected, uroplakin-3 (green) shows control ureters have stratified by this timepoint but *Exoc5* CKO ureters have not. (I) Higher magnification of the accumulation of LC3 in the E17.5 *Exoc5* CKO ureter. Statistical significance denoted by n.s.= not significant p ≥0.05, p* ≤ 0.05, p** ≤ 0.01 p*** ≤ 0.001. Scale bars = 20 µM.

### LC3 accumulates in *Exoc5* CKO urothelium

KEGG analysis of the most downregulated pathways revealed several metabolic mechanisms as being significantly perturbed following *Exoc5* ablation (Figure 2G). Given that the exocyst has been implicated in the initiation process of autophagic stress response (Bodemann *et al*., 2011), we reasoned that exocyst-mediated autophagy may be deficient in the urothelium and contributing to the downregulation of these metabolic pathways. Since LC3 is known to accumulate when autophagy is impaired (Runwal *et al*, 2019), we tested this by performing immunohistochemistry of the LC3 protein in E17.5 *Exoc5* CKO ureters. While control ureters showed no obvious abnormalities and only low levels of LC3, *Exoc5* CKO ureters displayed a strong accumulation of LC3 in the urothelium while the surrounding smooth muscle layer did not show significant increase (Figure 2H). It was noted that the *Exoc5* CKO urothelium showed LC3 puncta accumulation (Figure I, arrow), which is commonly observed when LC3 is not degraded from active autophagy.

### Inhibiting exocyst function causes impaired autophagy in urothelial cells

Though interactions between exocyst and autophagy related genes (ATG) have been previously reported, the functional consequence of perturbing this relationship in urothelial cells remains unclear. Here, we used 100µM endosidin-2 (ES2) to inhibit exocyst function (Zhang *et al*, 2016). ES2 treatment of primary human urothelial (pHUC) cells resulted in progressive vesicle accumulation over 24 hours, which was visible by phase contrast microscopy (Figure 3A,B). To determine if ES2 vesicle accumulation was the result of impaired autophagy in pHUCs, we performed western blotting of 24h ES2-treated pHUCs to measure protein levels of classic autophagy markers. Here, we measured a decrease in ATG5 levels, and significant increase in LC3II/I ratio and p62 accumulation after ES2 treatment (Figure 3C, D). In complement to the shift in LC3II/I ratio we also observed an increase in LC3 punta (Figure 3E). To test if the exocyst and ATGs directly associated in human urothelial cells, we performed immunoprecipitation of EXOC4 followed by western blotting for ATG7 using SV-HUC-1 immortalized urothelial cells and observed positive pulldown (Figure 3F). These data show that disrupting exocyst trafficking with ES2 treatment disrupts the process of autophagy.

**Figure 3.**
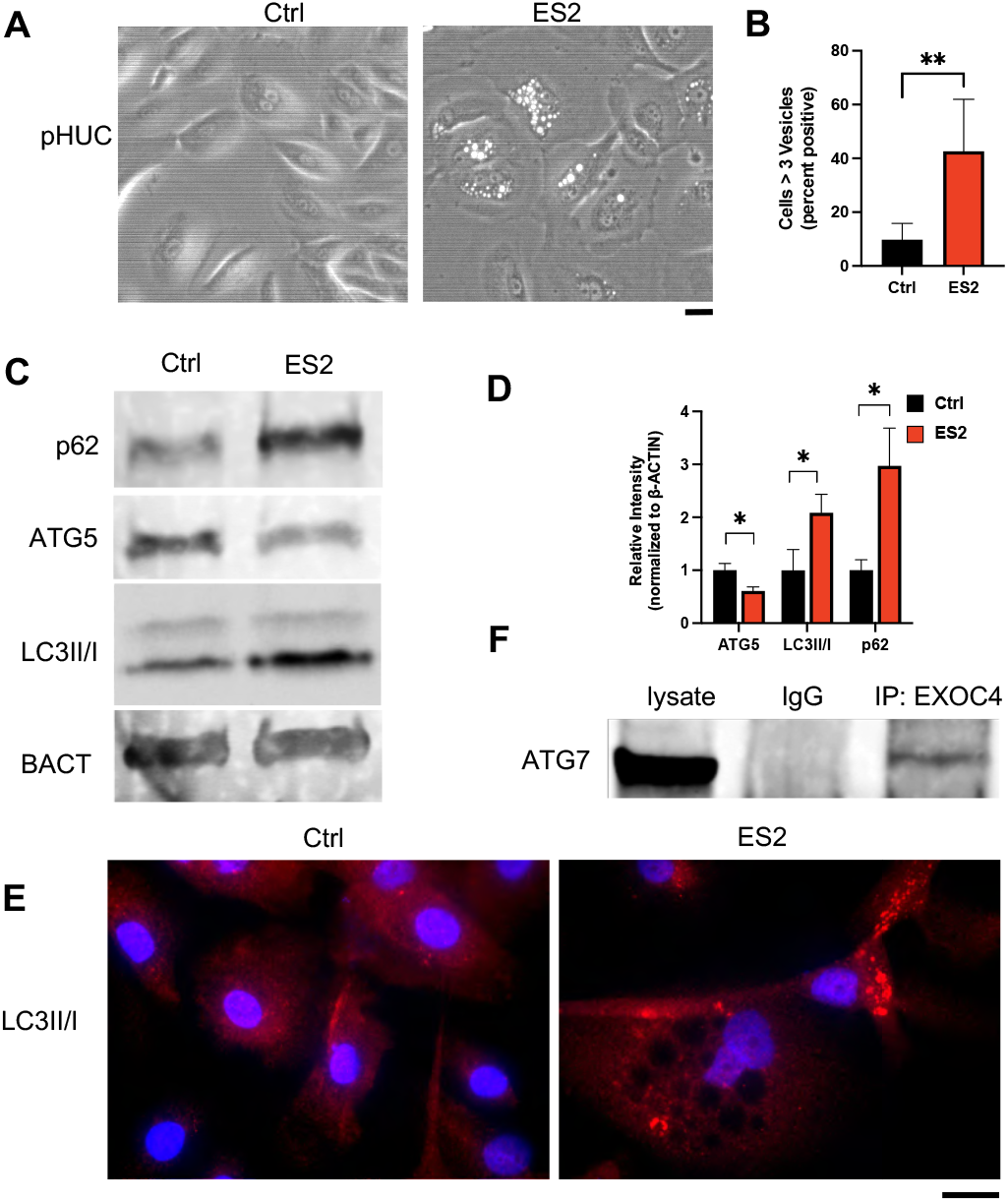
Inhibiting exocyst function in pHUCs results in decreased autophagy and vesicle accumulation. (A) Phase contrast microscopy of 24h 100µM endosidin-2 (ES2) treated pHUC cells display vesicle accumulation. (B) Quantification of control and ES2 treated cells with >3 vesicles (n=854, 778) show ES2 treated fields more likely to display pHUCs with vesicle accumulation. (C) Western Blot of 24h ES2 treated pHUCs indicated increased p62 and LC3 II/I ratio, with a reduction in total ATG5 also observed. (D) Quantification of immunoblotted autophagy markers after 24h 100µM ES2 treatment demonstrate a reduction of total ATG5 and increases in p62 and LC3II/I ratio. (E) Immunofluorescent detection of LC3II/I in 24h 100µM ES2 treated pHUCs display punctae accumulation. (F) In SV-HUC-1 cells, co-immunoprecipitation of EXOC4 successfully pulled down ATG7 denoting a biochemical protein-protein interaction. Statistical significance denoted by n.s.= not significant p ≥0.05, p* ≤ 0.05, p** ≤ 0.01. Scale bars = 20 µM.

To further demonstrate that inhibiting exocyst function had a detrimental effect on autophagy in urothelial cells, we ablated *Exoc5* in the adult bladder urothelium by crossing *Exoc5*^*FL/FL*^ with *Upk3a-GCE* mice, which express both tamoxifen-activated Cre and GFP under the uroplakin-3a promoter (Honeycutt *et al*, 2015). In parallel, *tdTomato* Cre-reporter mice were crossed with *Upk3a-GCE* mice to assess Cre recombinase activity and specificity after tamoxifen treatments (Figure 4A). The *Exoc5*^*FL/FL*^*;Upk3a-GCE* mice were treated with tamoxifen at 6-8 weeks of age to generate induced urothelial *Exoc5* knockout (*Exoc5-iUKO)* mice, with control mice defined as *Exoc5*^*FL/FL*^ mice treated with an identical regimen of tamoxifen. The *Exoc5-iUKO* survived with no gross abnormalities until euthanized 4 weeks later for bladder histology and electron microscopy. Scanning electron microscopy on the *Exoc5-iUKO* luminal bladder surface did not reveal any noticeable abnormalities in uroplakin plaque structures, and H&E histology was unremarkable. However, transmission electron microscopy (TEM) on *Exoc5-iUKO* bladder sections revealed a significant buildup of lysosomes as marked by electron dense organelles, which was similar to observations found in age-related lysosomal disease models (Figure 4B) (Truschel *et al*, 2018). To test for impairments in autophagy in these *Exoc5-iUKO* urothelial cells, we performed immunofluorescent detection of LC3 on bladder sections and observed classical large LC3 puncta formation at a highly increased instance over those seen in controls (Figure 4C,D). Taken together, these data indicate that the exocyst biochemically interacts with ATGs in urothelial cells and that inhibiting exocyst function detrimentally affects autophagy in a tissue-specific manner.

**Figure 4.**
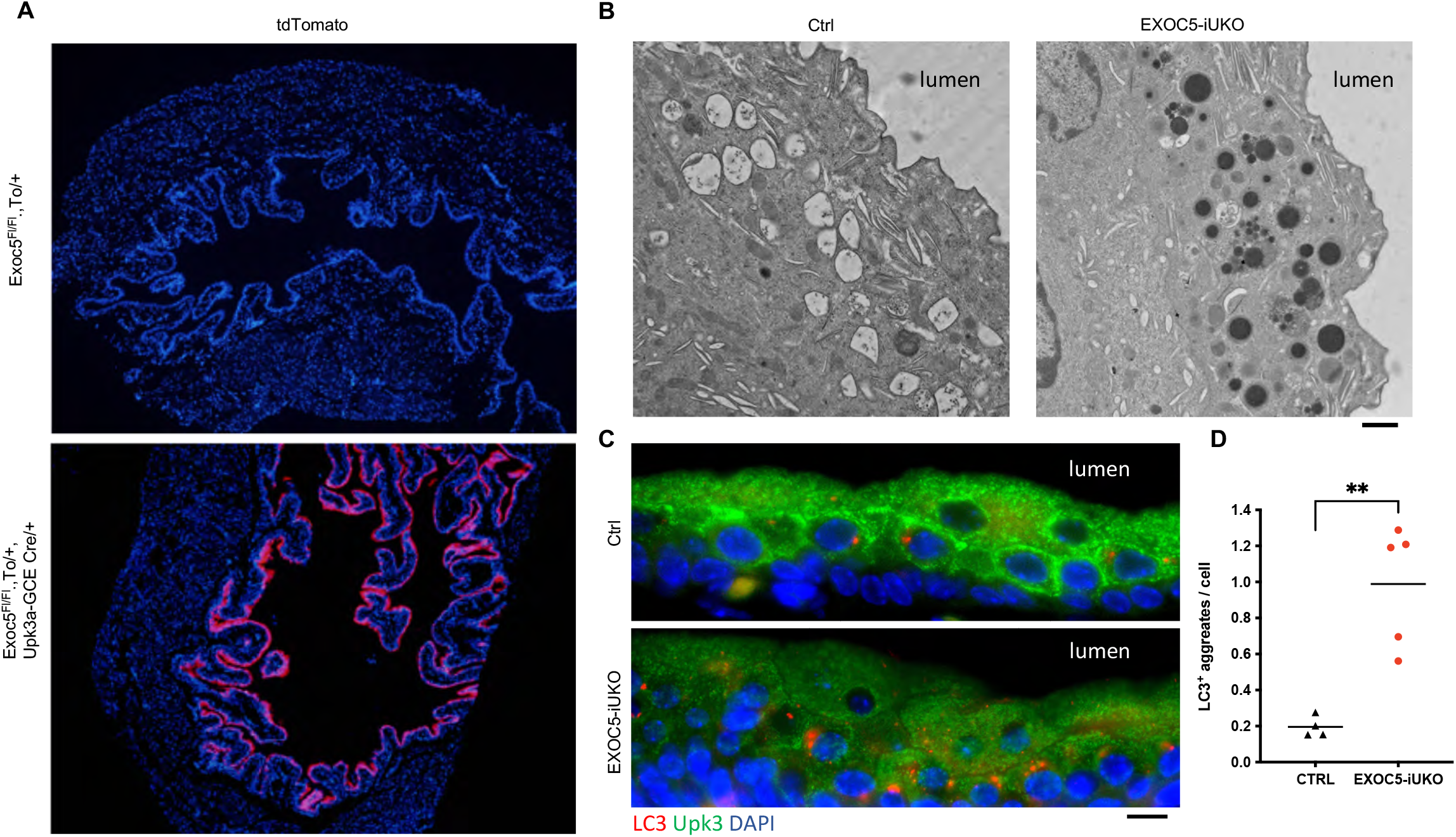
*Exoc5* ablation in adult bladder urothelial cells results in lysosomal accumulation. (A) Specificity of the Cre reporter system shown by tdTomato activation in superficial urothelial cells. (B) Transmission electron microscopy of bladder sections display lysosomal buildup as marked by electron dense organelles. (C) Immunohistochemistry of EXOC5-IUKO bladder sections displayed an accumulation of LC3+ puncta in the superficial urothelial cells. (D) Quantification of LC3+ puncta aggregates per cell with each dot representing a mouse. Statistical significance denoted by p** ≤ 0.01. Scale bars = 20 µM.

### Fn14 is upregulated by inhibiting autophagy in urothelial cells

We wanted to determine if there was a causal relationship between impaired autophagy and the noncanonical NF-κB signaling observed in our microarray data. To test this, we investigated whether Fn14 responded to disruptions of autophagy and other forms of stress in SV-HUC-1 cells. First, we treated SV-HUC-1 cells with 20µM cisplatin for 4 hours to assess the effect of cisplatin-induced DNA damage on Fn14 levels and found no significant induction of Fn14 (Figure 5A,B). Next, we exposed SV-HUC-1 cells to UV 40 J/m^2^ and found no significant Fn14 increase after 6 hours of recovery (Figure 5C,D). These results indicated that under two different forms of DNA damage induced cell stress, Fn14 was not upregulated in the measured timeframes. Next, to determine if exocyst depletion impacted SV-HUC-1 cells ability to respond to stress we generated shExoc5 stable knockdowns. We observed by western blotting that treating SV-HUC-1 cells with 200nM of autophagy inhibitor BafA1 for 24h caused a significant increase in Fn14 and that this response appeared synergistic in shExoc5 knockdown cells (Figure 5 E,F,G). 200nM BafA1 treatment of wild type SV-HUC-1 cells over a 5 day period greatly reduced colony formation (Figure 5H,I) with significantly less cell viablity observed after 3 days (Figure 5J). Interestingly, inhibiting autophagy with either BafA1 or VPS34 inhibitor (VPS34i) both induced vesicle accumulation that was similar to ES2-treated cells (Figure EV5). These results indicated that inhibiting autophagy with BafA1 was sufficient to induce Fn14 and promote cell death. These data suggest that EXOC5 knockdown added an additional stress in complement to the direct autophagy inhibition.

**Figure 5.**
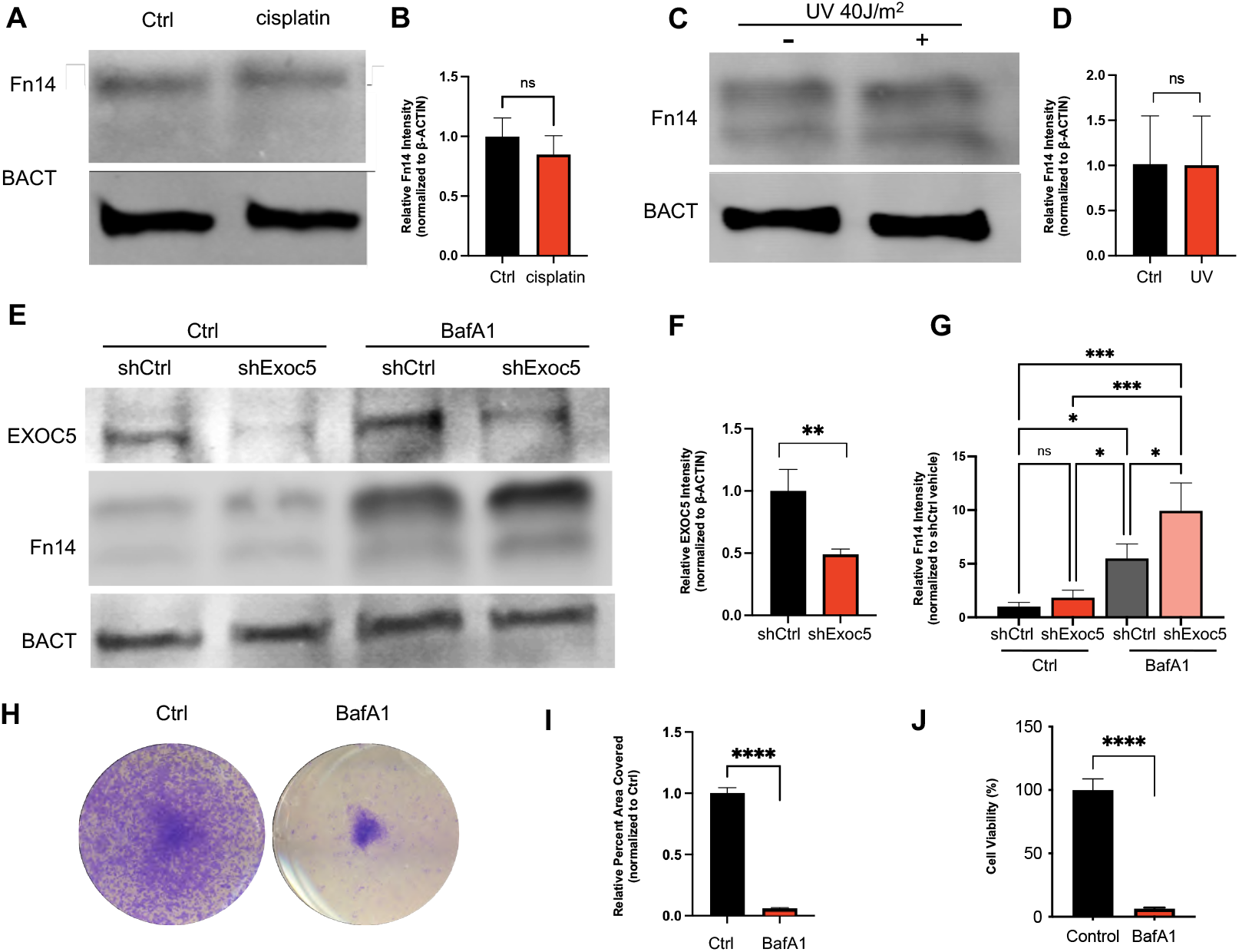
Autophagy inhibition induces NF-kB activator, Fn14, response. (A, B) Western Blot analysis of SV-HUC-1 cells treated with 20µM cisplatin for 4 hours to induce DNA damage showed no change in Fn14. (C, D) Western Blot analysis of SV-HUC-1 cells dosed with UV 40 J/m^2^ then left to recover for 6 hours showed no increase in Fn14. (E, F) Stably selected shExoc5 SV-HUC-1 cells displayed a synergistic sensitivity in inducing Fn14 response when treated with 200nM BafA1 to inhibit autophagy for 24h when compared to shCtrl. (G) Western blot quantitation of 200nM BafA1 treatment for 24h increases Fn14 in both control and shExoc5 groups and that this effect is highest in shExoc5 + BafA1 (H) SV-HUC-1 cells grown for a week in the presence of 200nM BafA1 then stained with crystal violet (CV) staining solution showed inhibiting autophagy significantly impaired growth. (I) Quantification of surface area coverage by CV staining. (J) CellTiter-Glo ATP-based cytotoxicity assay of SV-HUC-1 cells treated with BafA1 demonstrated cell death occurring by 72 hours. Statistical significance denoted by n.s.= not significant p ≥0.05, p* ≤ 0.05, p** ≤ 0.01 p*** ≤ 0.001.

### Inhibiting autophagy activates two waves of NF-κB signaling

Next, we wanted to determine if NF-κB signaling was responding to cell stress triggered by impaired autophagy in these urothelial cells. First, we performed immunocytochemistry on SV-HUC-1 cells treated with either 200nM BafA1 or 50µM VPS34i to determine if canonical or non-canonical NF-κB were being stimulated with autophagy inhibition. We measured the percentage of cells showing nuclear translocation of RelA (p65) as early as 6 hours after either BafA1 or VPS34i treatment (Figure 6A, Figure EV5), indicating that canonical NF-κB signaling was active. Nuclear translocation of RelA remained present through 48 hours, although varying levels of saturation were observed. To determine if non-canonical NF-κB was also being stimulated by autophagy inhibition, we performed a similar timecourse and measured the percentage of cells with p52 nuclear translocation. While p52 nuclear translocation was not significantly detected 6 hours, we found an increasing prevalence as time passed from 24 hours to 48 hours nearing the time of significant BafA1-induced cell death (Figure 6B). These data indicate that inhibiting autophagy results in canonical NF-κB activity and that prolonged treatment promotes non-canonical NF-κB activity.

**Figure 6.**
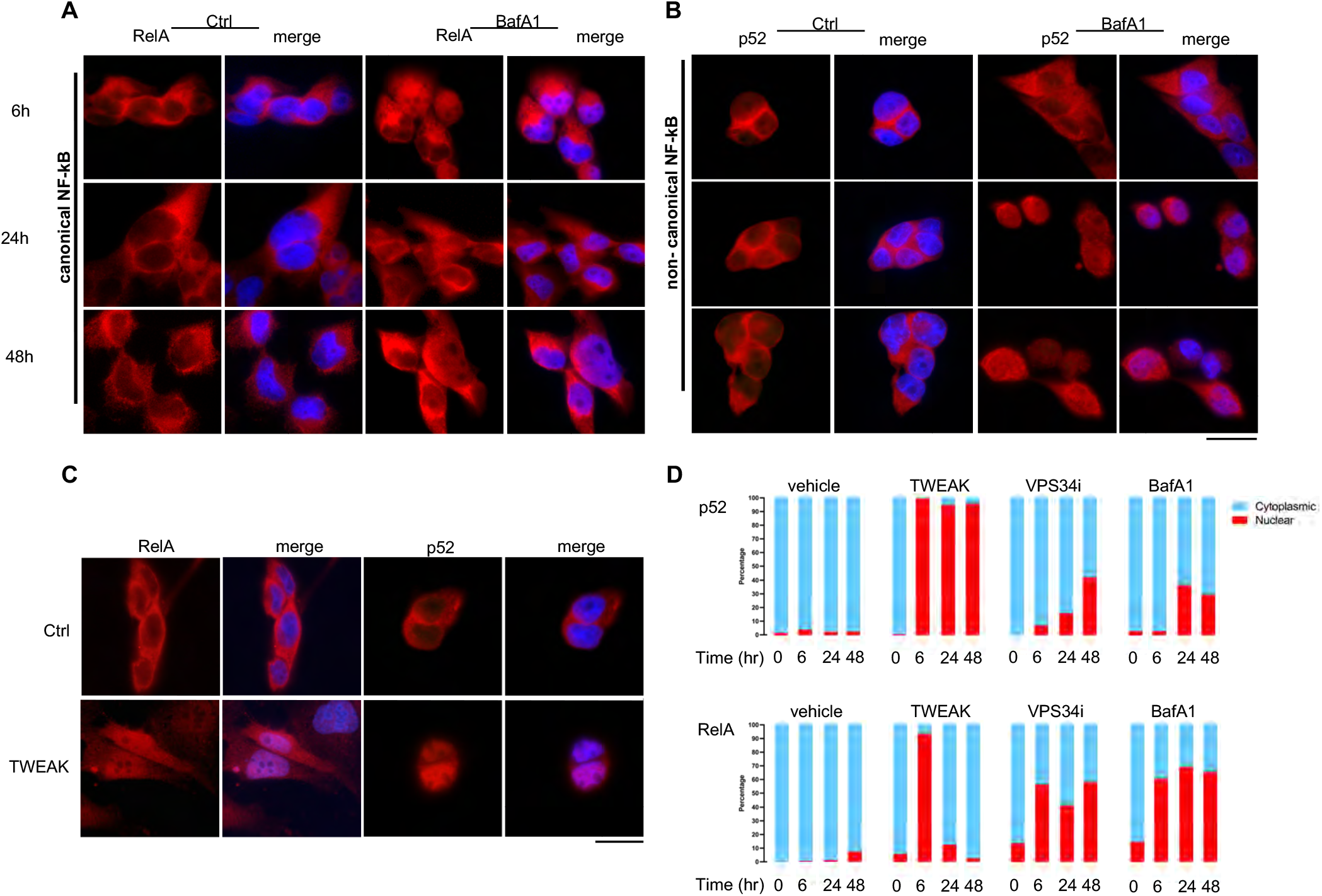
Inhibiting autophagy in SV-HUC-1 cells activates progressive canonical then non-canonical NF-kB signaling. (A) Immunocytochemistry of 200nM BafA1 treated SV-HUC-1 cells stained for RelA/p65 (red) showed nuclear translocation as early as 6h, with sustained localization observed at 24h and 48h. (B) Immunocytochemistry of 200nM BafA1 treated SV-HUC-1 cells demonstrated that nuclear translocation of non-canonical NF-kB constituent p52 (red) was not observed at 6h but appeared later at 24h and 48h. (C) TWEAK treated SV-HUC-1 cells stained for RelA/p65 (red) and p52 (red) demonstrated both canonical and noncanonical NF-kB signaling were active at 6h. (D) Quantification of 6h, 24h, and 48h time course detecting nuclear translocation of RelA/p65 and p52 in response to TWEAK, VPS34i, or BafA1.

While TNF superfamily receptors can activate canonical NF-κB, Fn14 is one of the few known receptors to primarily activate non-canonical NF-κB signaling. To confirm that Fn14 signaling was active in SV-HUC-1 cells, we applied its only known ligand, the cytokine TWEAK, and observed a strong increase of both p52 and RelA nuclear translocation by 6 hours (Figure 6C). However, only p52 remained at significant levels by 24 and 48 hours (Figure 6D). This indicated non-canonical NF-κB signaling can be activated in human urothelial cells through the TWEAK-Fn14 signaling axis and that while both NF-κB pathways can be activated by TWEAK in urothelial cells, the promotion of non-canonical NF-κB signaling was sustained for longer periods. Taken together, these data indicate TWEAK-Fn14 activate canonical NF-κB in response to autophagy inhibition as an initial response to stress but that a second wave of noncanonical NF-κB follows if the insult persists (Figure 6D).

### z-VAD-FMK rescues cell death and ureter obstruction in *Exoc5* CKO ureters

Since TWEAK-Fn14 and non-canonical NF-κB signaling can initiate caspase activity and cell death, we explored whether *Exoc5* CKO urothelial death could be prevented *in vivo* with a pan-caspase inhibitor (Ikner & Ashkenazi, 2011; Martin-Sanchez *et al*, 2018; Vince *et al*, 2008). To perform this, we administered 5 μg/g z-VAD-FMK by intraperitoneal injection to timed mated female mice at E16.5 and collected embryos at E18.5. Under normal conditions, the *Exoc5* CKO urothelium fails to stratify and undergoes cell death between E16.5 and E17.5 with full ureter obstruction by E18.5 as the underlying mesenchyme obliterates the ureter lumen (Fogelgren *et al*., 2015; Lee *et al*., 2016) (Figure 7A). However, when administering z-VAD-FMK at E16.5, we observed no urothelial cell death and complete rescue of proper ureter formation (n=9/9 *Exoc5* CKO embryos) (Figure 7A). Furthermore, the *Exoc5* CKO urothelium successfully stratified as seen by UPK3 immunofluorescent signal in the umbrella cells (Figure 7A). These *in vivo* results agreed with our *ex vivo* explant data and previously published data (Lee *et al*., 2016), which indicated that obstruction was the subsequent result of widespread urothelial cell death. With the obstruction prevented, we observed that hydronephrosis was avoided in these *Exoc5* CKO embryos as well (Figure 7B). In addition, while *Exoc5* CKO mice typically die 8-14 hours after birth E16.5 rescue trials revealed that all z-VAD-FMK-treated Exoc5 CKO mice successfully survived to adulthood.

**Figure 7.**
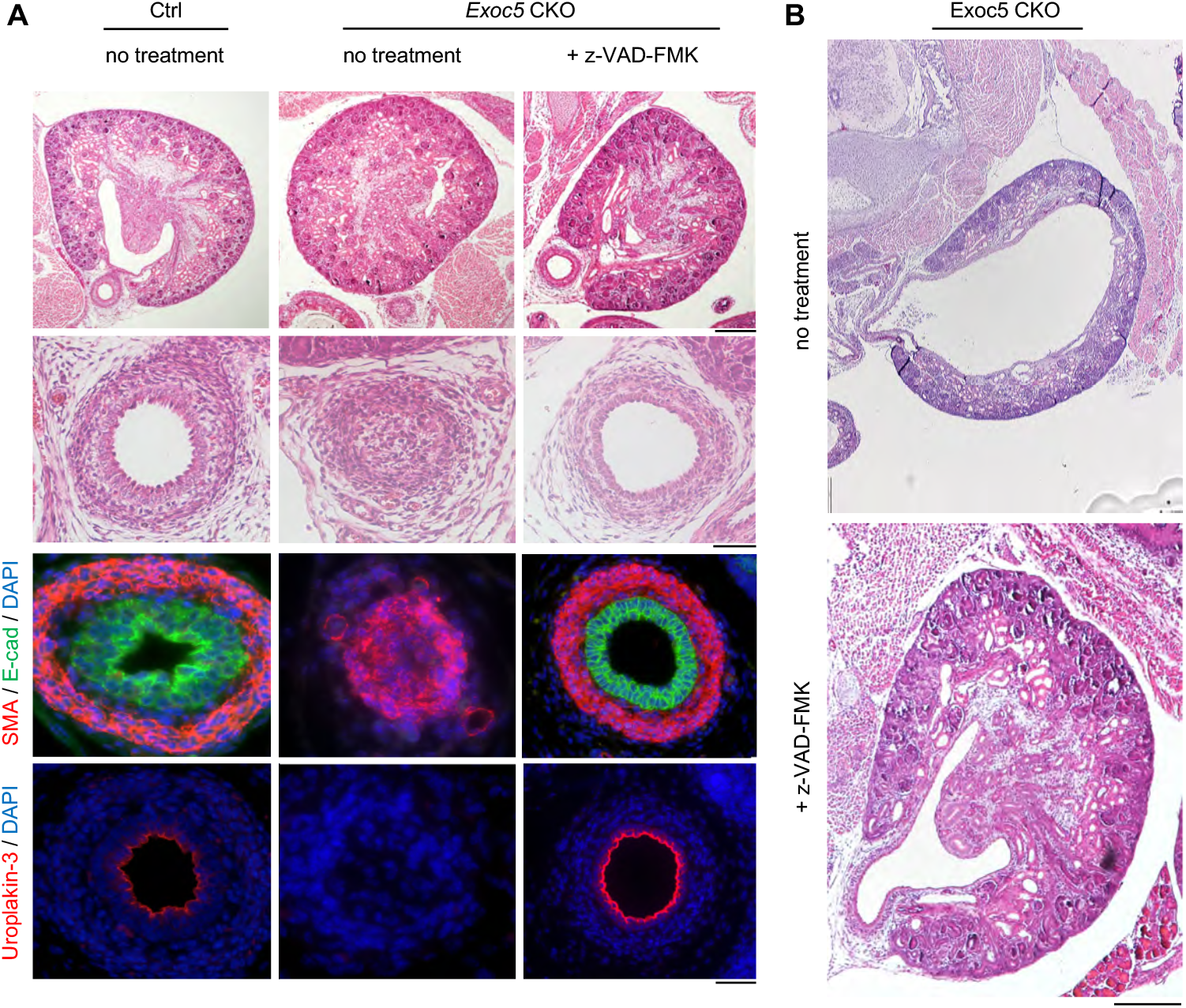
Single dose of pan-caspase inhibitor z-VAD-FMK at E16.5 rescues ureter obstruction in *Exoc5* CKO embryos. (A) H&E staining of E18.5 embryonic ureters from control, *Exoc5* CKO with no treatment, and *Exoc5* CKO with z-VAD-FMK ureters demonstrate rescue of ureter obstruction and the absence of cell death (Row 1 low mag., Row 2 high mag.). Immunohistochemistry staining of the same series showed *Exoc5* CKO mice treated with z-VAD-FMK rescued stratification as indicated by E-cadherin (green) and SMA (red). Positive uroplakin-3 (UPK3) (red) staining in z-VAD-FMK treated *Exoc5* CKO indicated successful differentiation of superficial cells. (Row 1 scale bar = 200 μm, Rows 2-4 scale bar = 20 μm). (B) H&E staining of E18.5 kidneys from *Exoc5* CKO mice with and without z-VAD-FMK treatment showed subsequent absence of hydronephrosis (scale bar = 200 μm).

## Discussion

While defects in membrane trafficking are well known to contribute to a breadth of human diseases it remains unclear whether or not aberrant exocyst function contributes to developmental defects of the urinary tract, such as COU (Coulter *et al*, 2020; Uhm *et al*, 2017; Van Bergen *et al*, 2020). The phenotypic mirroring of human COU by the *Exoc5* CKO mouse offers a congenital model for deconstructing the molecular mechanisms underlying the onset of obstructive uropathy. Nephrogenesis is dependent on ureteric bud branching into the metanephric mesenchyme, which starts around E11.5, and we have previously shown that Cre recombinase driven by the *Ksp*-cadherin promoter is active by this time (Lee *et al*., 2016). Thus, *Exoc5* is not required for branching morphogenesis by the ureteric bud, and the urinary tract appears morphologically normal until E17.5. However, we observed that *Exoc5* ablation in the urothelium resulted in autophagy impairment and non-canonical NF-κB activity prior to cell death at E17.5, where we further report TUNEL-positive and cleaved caspase 3-positive signal only from a small pool of cells that have already detached from the ureter wall. These data suggest the loss of cell anchorage in a minority of urothelial cells might play a role in inducing anoikis, which is a form of apoptosis. Taken together, these data support a model where exocyst depletion promotes impaired autophagy, and that autophagic deficiencies in turn activate progressive NF-κB signaling and eventually cell death. While the exact series of mechanisms will need to be further investigated, it is clear that the Exoc5 CKO model of COU can be rescued with the pan-caspase inhibitor z-VAD-FMK.

The relationship between defective autophagy and NF-κB signaling is intricately tied to cellular stress and damage response. For example, deficiencies in autophagy have been shown to activate canonical NF-κB in part through an accumulation of p62, which serves as a scaffold for the TRAF6 and RIP1 complex that drives canonical NF-κB activation (Meng & Cai, 2011; Wooten *et al*, 2005). Autophagy typically suppresses p62 accumulation, thereby guarding against unnecessary canonical NF-κB signaling (Mathew *et al*, 2009). In the data presented here, we observed LC3 and p62 accumulation when inhibiting exocyst function, which suggests a path for activating canonical NF-κB as part of a survival response in urothelial cells. More specifically, it is reasonable to posit that inhibiting EXOC5 would further impact the formation of the RALB-EXOC84 effector complex which would have a direct impact on the ability to initiate autophagic response to stress. The relationship between autophagy deficiency and canonical NF-κB activation is further supported here by BafA1 treatment induced RelA nuclear translocation. The initial canonical NF-κB response to BafA1 treatment may be an attempt to promote cell survival in the face of cell stress. However, these data further indicated that sustained stress in urothelial cells promotes a slower second non-canonical NF-κB response.

One way to detect defective autophagy is the accumulation of LC3, as was observed in *Exoc5* CKO ureter urothelium at E17.5. This LC3 accumulation was enhanced in *Exoc5* ablation in adult bladder urothelium using *Upk3a-GCE* mice. The urothelial *Exoc5* ablation presented here in the ureter and bladder highlight a tissue and time specific relationship between the exocyst and autophagy. It is likely that *Exoc5* ablation in the bladder urothelium presented with larger LC3 punctae because there was a longer time frame (four weeks) for this buildup to occur. This observation would also support the hypothesis that an exocyst-mediated event initiated the stress response in the developing ureter and that the impaired autophagy contributed to NF-κB activation.

Interestingly, ablation of *Exoc5* would also impact formation of the RALB-EXOC2 effector complex which directly activates TBK1. TBK1 plays a critical role in inhibiting cell death and maintaining non-canonical NF-κB by phosphorylating NF-κB-inducing kinase (NIK), which leads to NIK degradation (Chien *et al*., 2006; Jin *et al*, 2012). NIK degradation is necessary for preventing p100 processing to p52 (Xiao *et al*, 2001). By extension, the urothelial stress induced by *Exoc5* ablation could reasonably be expected to impact RALB-EXOC2 effector complex interaction with TBK1 and thereby promote non-canonical NF-κB signaling through NIK accumulation. Since, the exocyst acts as a moderator of these pathways, it will be necessary in the future to confirm these interactions in ureter development.

Our observation that non-canonical p52 nuclear translocation acts as a slower response complements results reported by Saitoh et al. 2003, in which TWEAK was directly used to stimulate non-canonical NF-κB independent of TNFα in mouse embryonic fibroblasts (MEFs) and found p52 nuclear translocation was delayed in response when compared to canonical RelA nuclear translocation (Saitoh *et al*, 2003). Under these conditions, canonical RelA responded first but was followed by non-canonical p52 after 8 hours, whereafter p52 remained in the nucleus for at least 24 hours (Saitoh *et al*., 2003). Depletion of exocyst function and subsequent autophagic deficiency contribute added complexity to the cells ability to deal with stress. Progression to TWEAK-Fn14-mediated non-canonical NF-κB activation is intriguing because TWEAK can induce several cell death pathways including caspase-mediated death but to date there is not yet a strong body evidence connecting this pathway to anoikis (Chicheportiche *et al*, 1997; Nakayama *et al*, 2003). Here, it is important to state that these data are not suggesting Fn14-mediated non-canonical NF-κB is singularly dependent on the effects of autophagic deficiency, nor that this is the sole driving factor behind *Exoc5* CKO ureter cell death, but rather these data indicate a role for impaired autophagic flux in promoting a sustained TWEAK and Fn14 environment. More significantly, we report that the pan-caspase inhibitor z-VAD-FMK can prevent urothelial cell death and ureter oblation from occurring in the Exoc5 CKO mouse model.

In summary, these data demonstrate that urothelial cell death is initiated independent of aberrant urothelial stratification and is the underlying pathological driver of congenital obstructive uropathy in the *Exoc5* CKO mouse model. We further implicate the effects of impaired autophagy in responding to stress events during ureter development, whereby there is progressive promotion of canonical and then non-canonical NF-κB signaling. These findings may provide insight into the pathological events contributing to human COU.

## Methods

### Animals

All animal procedures and protocols were conducted in accordance with IACUC specifications approved by the University of Hawai’i Animal and Veterinary Services. Dr. Fogelgren’s IACUC approved protocol is #11-1094 and the University of Hawaii has an Animal Welfare Assurance on file with the Office of Laboratory Animal Welfare (OLAW), assurance number is A3423-01. Mice were housed under standard conditions with a 12-hour light cycle with water and food *ad libitum*. The floxed EXOC5 mouse strain (EXOC5^FL^) was generated and used as previously described in Fogelgren, et. al, 2015, as was the *tdTomato* Cre reporter strain (Jackson Laboratories stock #007909). The Ksp-Cre and UPK3a-GCE mouse strains were obtained from Jackson Laboratories (stock # 012237 and 015855, respectively). All mice were on a C57/Bl6/J inbred background. For timed matings, females mated with a male were monitored for abdominal bulging beginning fourteen days after arranging timed mating. Embryos were dissected in the morning and measured by embryonic body length and staged using Theiler staging criteria (TS) to ensure the developmental stage of each embryo. 6-8 week old *UPK3a-GCE* mice were fed tamoxifen-containing chow (Cat #TD.130860, ENVIGO) for two weeks, and then after four weeks with normal chow, bladders were collected for histology analysis. For rescue experiments, timed mated female mice were i.p. injected at E16.5 with z-VAD-fmk (R&D Systems, Cat #FMK001) at 5 μg/g body weight in 10% DMSO in PBS. Embryos were collected 48 hours after injection, with caudal torsos collected for histological analysis and DNA isolated for genotyping.

### Histology and immunohistochemistry

Caudal torsos of *Exoc5* knockout and control animals were dissected and fixed in 4% formaldehyde overnight with rocking at 4 °C. Samples were embedded in paraffin according to standard protocols and cut into 5 μm sections. Staining and immunohistochemistry procedures were performed as previously reported (Fogelgren *et al*., 2015). Primary antibodies used were: anti-E-cadherin at 1:200 (Cell Signaling, Cat #3195), anti-Smooth Muscle Actin at 1:800 (Millipore Sigma, Cat #A2547), anti-Uroplakin-3 at 1:100 (ARP, 03-610108), and anti-LC3 at 1:500 (Cell Signaling, Cat. #12741). Secondary antibodies (Dylight, 35552 and 35560) and were used at 1:800 at wavelengths 488 and 594 nm. Stained sections were analyzed using a fluorescent Olympus BX41 microscope. LC3 quantification analysis was performed using ImageJ software (NIH).

### Co-immunoprecipitations (co-IPs) and western blots

Samples were lysed in co-immunoprecipitation buffer (50nM Tris-HCL (pH 8), 150mM NaCl, 5mM EDTA, 0.5% NP-40, 1mM DTT, 20mM NaF) containing phosphatase and protease inhibitors via mechanical homogenizing and vortex. Samples were placed microcentrifuge and spun at 14,000 rpm at 4°C for 30 minutes. The supernatant was removed and proteins were quantified by Bradford’s assay. Protein input for co-immunoprecipitation was 2 mg and 8 μg of antibody was used per 2 mg of protein input. Protein samples were incubated with antibody against EXOC4 (Enzo, ADI-VAM-SV016-D) or rat IgG control (Fisher, Cat #02-9602) overnight with end-to-end rotation in 4°C. Immune complexes were then pulled down using Protein A/G Magnetic Beads (Thermo Fisher, Cat #88802), boiled in 2x Laemmli Sample Buffer (Biorad, Cat #1610737) with β-mercaptoethanol. The supernatant was run using SDS-PAGE and transferred onto nitrocellulose membrane using Trans-Blot Turbo Transfer System. The membrane was blocked in 5% nonfat milk for 1 hour and probed with primary antibody overnight. Secondary antibodies at 600 nm or 800 nm (Licor IRDye) were incubated at 1:10,000 for 1 hour followed by 3x PBSt washes and scanned on Odyssey CLx Imaging System.

Other antibodies used in western analysis include p62 (Cell Signaling, Cat #16177S), ATG7 (Cell Signaling, Cat #8558), ATG5 (Cell Signaling, Cat #12994), EXOC5 (SEC10) (Santa Cruz, Cat #sc-514802), LC3 (Cell Signaling, Cat #12741), Fn14 (Abcam, Cat #EPR3179). ImageStudioLite (LiCor) was used to analyze and pseudocolor all scans to grayscale.

### Cell culture

For immunofluorescence, cells were seeded on coverslips and grown overnight. Treatment with 100µM endosidin-2 (Cayman, 21888) (Zhang C et al, 2016; Fujimoto et *al*, 2019; Leskova et al, 2020) 50µM VPS34i (Cayman, 17392), 200nM BafA1 (Sigma-Aldrich, B1793) were applied in fresh media change at implicated time and grown under standard conditions at 37°C and 5% CO_2_ for the respective timepoints listed. Cells were washed three times with PBS followed by fixation for 10 minutes in 4% PFA. Cells were permeabilized for 10 minutes with 0.1% Triton X-100 with 3x PBSt washes occurring between each step. Cells were blocked in 5% BSA in PBSt for 1 hour then incubated with the respective primary antibody (1:100) overnight at 4°C. Primary antibodies used were EXOC5 (SEC10), ATG5, p65 (Cell Signaling, Cat# 8242S, p52 (Millipore, Cat# 05-361), GOL97 (Santa Cruz, Cat #sc-59820), Fn14. Cells were washed 3x PBSt then incubated with secondary antibody (Dylight) for 1 hour, washed with 3x PBSt for 5 minutes and mounted using VECTASHIELD® Antifade Mounting Medium.

### Crystal violet staining

300k SV-HUC-1 cells were seeded and transfected with 3µg MISSON pLKO.1-puro non-mammalian targeting control shRNA (Sigma, Cat# SHC002) or shExoc5 (Sigma, Cat# TRCN0000061963) using RNAiMAX (Thermo Fisher, Cat #13778075,) according to the manufacturer’s protocol. Transfected cells were grown for 72 hours followed by puromycin selection. Stable cell lines were confirmed for EXOC5 knockdown by western and plated at 100k cells per well in 6 well plates in triplicate and grown for 1 week under vehicle control or 200nM BafA1 followed by staining with 0.1% crystal violet (CV) solution. CV solution was removed with repeated H_2_O washes and the plate was scanned. Percent area coverage was measured using ImageJ software and set relative to control.

### Cell Viability Assay

10K SV-HUC-1 cells were seeded per well of a 96-well opaque plate. Cells were treated and then subsequently assessed for their cell viability by the CellTiter-Glo Luminescent Cell Viability Assay (Promega, Cat #G7570). The assay measures ATP as a biomarker of metabolically active cells by the luminescence released from the conversion of Beetle Luciferin + ATP with the enzyme catalyst Ultra-Glo recombinant luciferase and Mg_2+_ to oxyluciferin + AMP + PPi + CO2 and light. 50 µL of CellTiterGlo reagent and buffer mix was added to 50 µL of cell media in each well and luminescence was measured by SpectraMax M3 (Molecular Devices,).

### *Ex vivo* ureter culture model

Timed matings were set up and embryos were collected at gestational age 15.5 (E15.5). Ureters were microdissected and placed on 1.0μm sterile filters at the air-liquid interface in a 12 well plate in 50/50 DMEM/HAMS F12 media supplemented with 5μg/ml transferrin, 100μg/ml penicillin, 100U/ml streptomycin. The ureters were cultured for 72 hours at 37°C. Gross images and peristalsis videos were taken using an Olympus CKX41 microscope. For samples that were treated with chemical agonists or antagonists, media containing the compounds were changed daily; and samples were cultured for 72 hours at 37°C. After 72 hours in culture, the ureter explants were fixed with 4% paraformaldehyde overnight and subsequently placed in sucrose for cryo-sectioning and histological analysis. Alternatively, the explants were collected for RNA extraction.

### Affymetrix Clariom D GeneChip microarrays

Ureters were microdissected from E16.5 embryos (n=3 *Exoc5*^*FL/FL*^ control and n=3 *Exoc5*^*FL/FL*^*;Cre*^*Ksp*^ CKOs) and RNA was isolated as previous described (Lee *et al*., 2016). After confirming RNA quality with an Agilent Bioanalyzer, gene expression profiling was performed at the University of Hawaii Genomics and Bioinformatics Shared Resources using Affymetrix Clariom D gene chips. Transcriptome analysis software (TAC, Affymetrix) was used to analyze and identify differential expression between wild type and Exoc5-CKO samples. GraphPad Prism 9 was used to generate a volcano plot and KEGG analysis was performed with the Database for Annotation, Visualization and Integrated Discovery (DAVID) (Huang *et al*, 2009; Sherman *et al*, 2008).

### Statistical analysis

All experiments were performed at least twice in triplicate with the most representative images shown. Error bars are presented as means ± SD and differences between groups were analyzed by two-tailed Student’s t test or one-way (ANOVA) where listed respectfully. Statistical significance was accepted at p* ≤ 0.05, p** ≤ 0.01 p*** ≤ 0.001 p**** ≤ 0.0001.

## ACKNOWLEDGMENTS

This work was supported by grants from the National Institutes of Health [grant numbers R01DK117308, R03DK100738, P20GM103456-06A1-8293 to B.F.]; and the March of Dimes [Basil O’Connor Starter Scholar Research Award, grant number #5-FY14-56 to B.F.]. We thank Tina Carvalho at the University of Hawaii Biological Electron Microscope Facility for her expertise in electron microscopy, the Histopathology Core (supported by NIH G12MD007601, P30GM103341, and U54MD007601) for outstanding histology services, and the Genomics and Bioinformatics Shared Resources for gene profiling support (P30CA071789).

---

**Expanded view 1.**
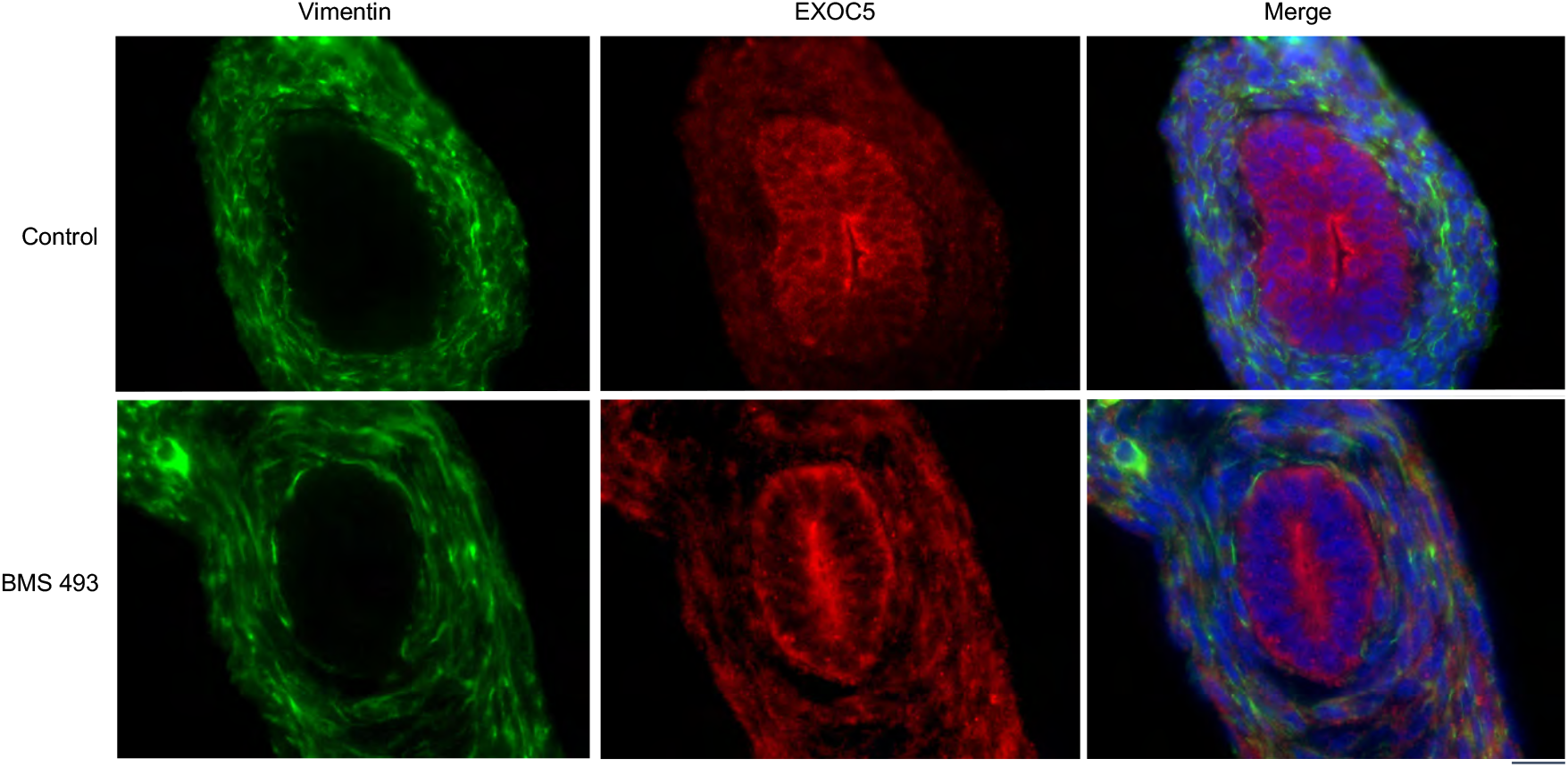
EXOC5 immunostaining in *ex vivo* cultured ureter explants treated with BMS493. Exoc5 is expressed throughout the urothelial cells with a concentration at the apical luminal membrane. No apparent difference in distribution is observed when urothelial differentiation is blocked with BMS493.

**Expanded view 2.**
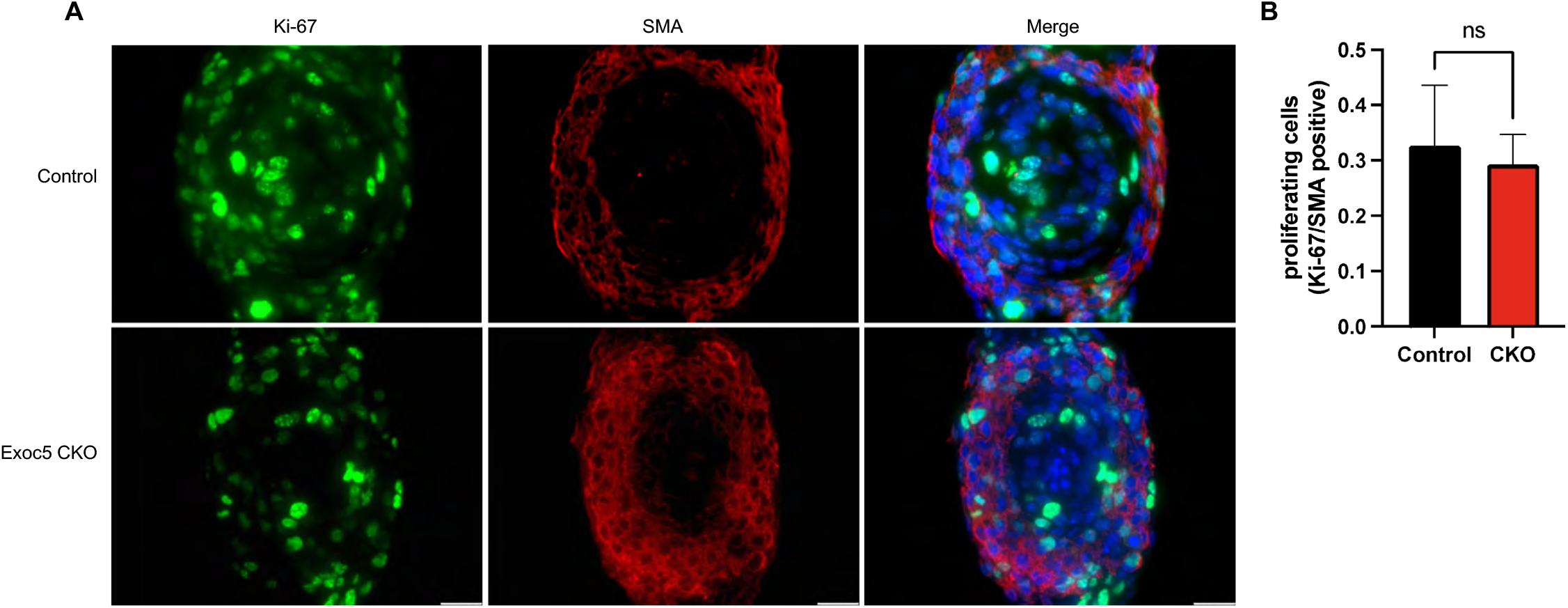
Immunostaining of Ki67 to measure proliferation rates in *Exoc5* CKO *ex vivo* cultured ureter explants. (A) Control and *Exoc5* CKO *ex vivo* cultured ureter explants displayed no significant differences in SMA distribution or Ki-67 abundance. (B) Quantitation of Ki-67/SMA show no significant differences after culturing for 72 h, indicating explants did not have the fibroproliferative response that was seen in vivo Exoc5 CKO ureters. n.s.= not significant p ≥0.05.

**Expanded view 3.**
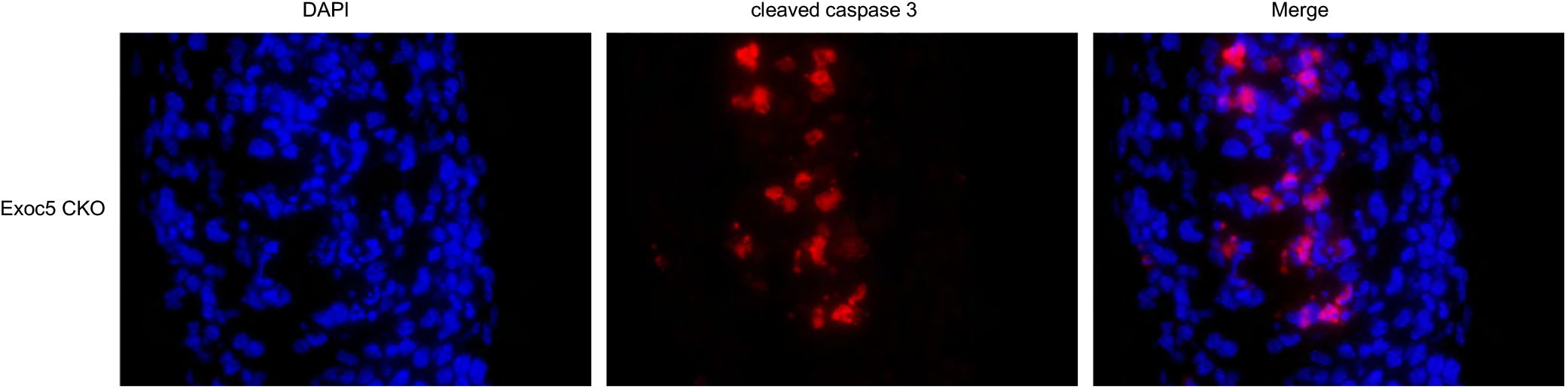
Immunostaining of cleaved caspase 3 in *Exoc5* CKO *ex vivo* cultured ureter explants. Cleaved caspase 3 is detected in a portion of cells that undergo basement membrane detachment and cell sloughing.

**Expanded view 4.**
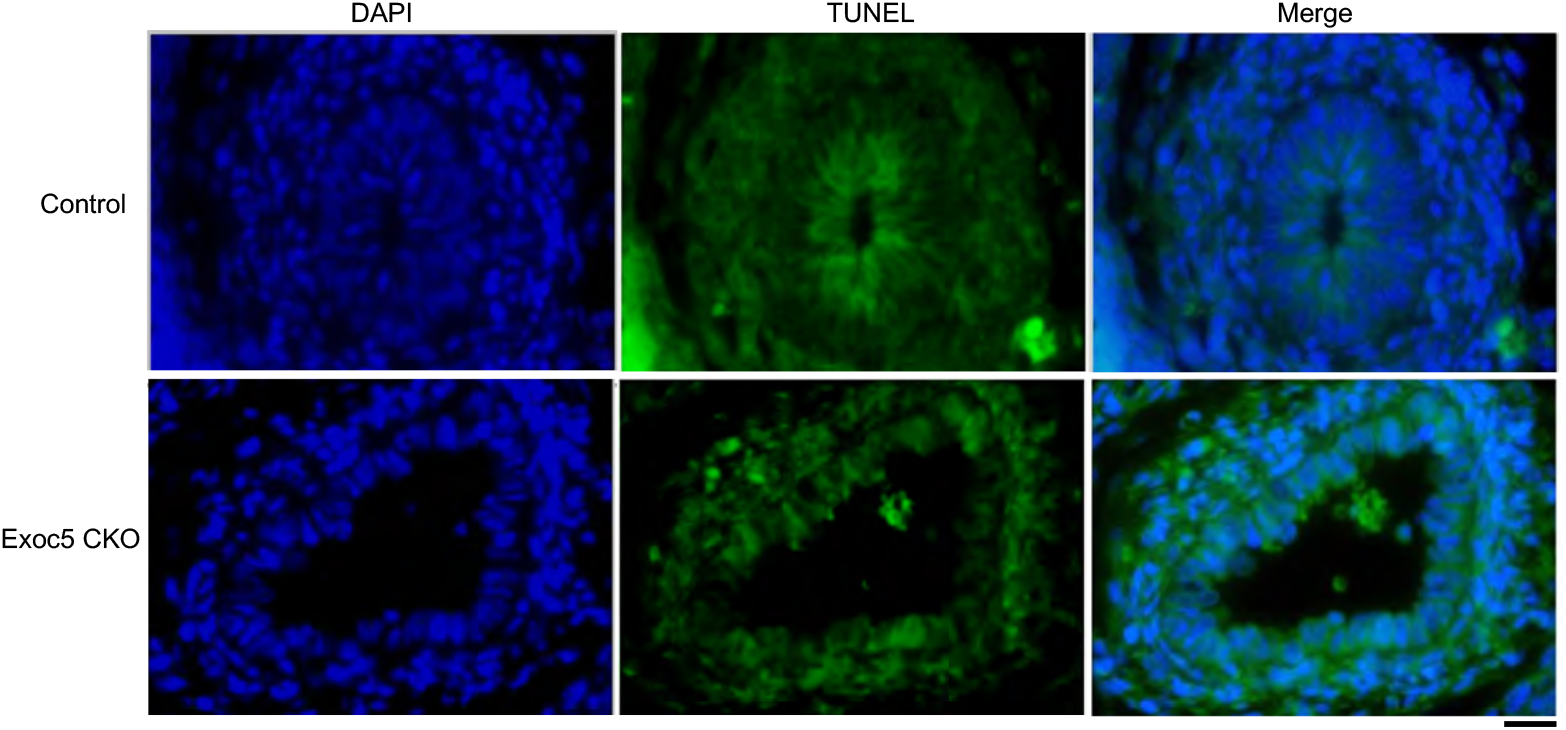
TUNEL staining of E17.5 *Exoc5* CKO ureters revealed few TUNEL-positive urothelial cells. Only detached urothelial cells sloughing off into the lumen of E17.5 CKO ureters were TUNEL-positive.

**Expanded view 5.**
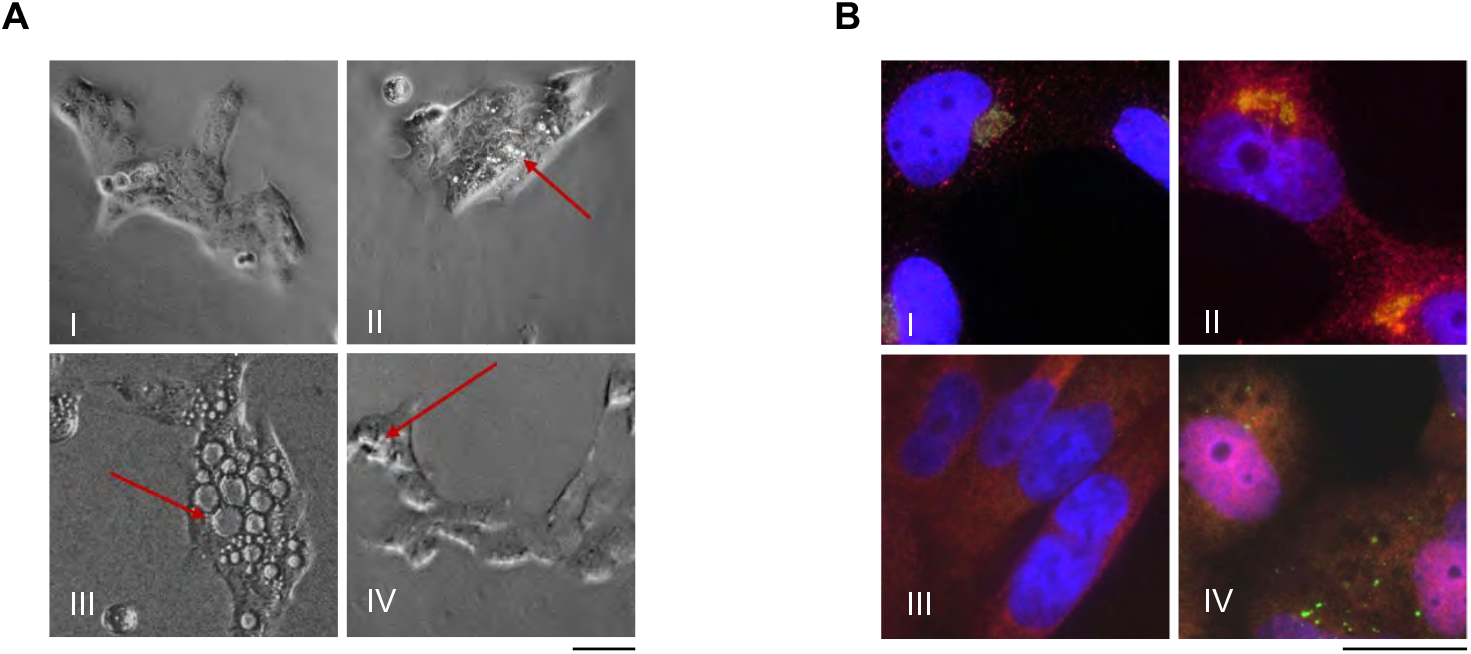
(A) Phase contrast microscopy of SV-HUC-1 cells treated with either (A-I) vehicle control, (A-II) 400 µM Endosidin-2, (A-III) 50 µM VPS34i, or (A-IV) 200 nM BafA1 for 24h. (B) Immunocytochemistry of SV-HUC-1 cells treated with vehicle control (I, III) or 50 µM VPS34i (II, IV) for 6h. (B-I,II) Fn14 (red) localization in relation to the Golgi complex as stained by Golgin97 (green). (B-III,IV) Immunohistochemistry of p62 (green) and nuclear translocation of RelA/p65 (red) in response to 50 µM VPS34i treatment. Scale bar = 20 µM

